# Complexity of spatiotemporal plantar pressure patterns during everyday behaviours

**DOI:** 10.1101/2023.01.27.525870

**Authors:** Luke D Cleland, Holly M Rowland, Claudia Mazzà, Hannes P Saal

## Abstract

The human foot sole is the primary interface with the external world during balance and walking, and also provides important tactile information on the state of contact. However, prior studies on plantar pressure have focused mostly on summary metrics such as overall force or centre of pressure under limited conditions. Here, we recorded spatiotemporal plantar pressure patterns with high spatial resolution while participants completed a wide range of daily activities, including balancing, locomotion, and jumping tasks. Contact area differed across task categories, but was only moderately correlated with the overall force experienced by the foot sole. The centre of pressure was often located outside the contact area or in locations experiencing relatively low pressure, and therefore a result of disparate contact regions spread widely across the foot. Non-negative matrix factorisation revealed low-dimensional spatial complexity that increased during interaction with unstable surfaces. Additionally, pressure patterns at the heel and metatarsals decomposed into separately located and robustly identifiable components, jointly capturing most variance in the signal. These results suggest optimal sensor placements to capture task-relevant spatial information and provide insight into how pressure varies spatially on the foot sole during a wide variety of natural behaviours.

## Introduction

The foot sole acts as an interface between our body and the environment, and its placement relative to the rest of the body determines stability in balance and gait. The foot sole is also a sensory organ innervated by thousands of tactile afferents that transmit information about dynamic contact parameters to the spinal cord and the brain. Indeed, after the hands and the face, the soles of the foot are the most heavily innervated regions of the body (1, 2). The importance of this sensory feedback is highlighted when sensation is impaired or lost, as seen in peripheral neuropathy or via experimental interventions, leading to increased sway velocity (3) and gait variability (4, 5), resulting in an increased risk of falls (6). The nature of tactile stimuli acting on the foot sole is very different from those encountered by our hands. For example, typical forces are much higher on the foot than on the hands, and in fact the foot sole is subjected to some of the highest loads of the entire body, experiencing over 2000 Newtons during jogging and jumping (7, 8). Surfaces that humans typically step on are also relatively flat, while we avoid surfaces with highly uneven height profiles in order to keep the body stable and prevent injury to the foot. Finally, in contrast to the hands, most of the interactions by the foot with the environment, at least in modern times and western societies, are mediated through a shoe (9), which acts as a barrier and transforms the pressure patterns acting on the foot sole (10). Together, these considerations imply a unique set of spatiotemporal pressure patterns that are typically experienced on the foot, but which are quite different from those on any other region of the body.

Previous investigations of the pressure patterns experienced on the foot sole have mostly focused on identifying differences in gait caused by various disorders and impairments, such as peripheral neuropathy. Existing research has tended to focus on static balance and normal gait (11–14). However, these activities do not cover all behaviours carried out in everyday life; we also rely on our feet during common activities such as stair climbing, jumping, and navigating unstable or uneven ground. While existing research has been valuable for our understanding of gait, a thorough characterisation of plantar pressure patterns from a sensory perspective is important for multiple reasons. First, to understand how a sense transduces and processes information, we need to characterize the range of natural sensory experiences that this sense is exposed to. Second, when studying a sense in carefully controlled conditions in a laboratory, we want to ensure that the artificial stimuli used fall within the range of those naturally occurring. Finally, when building devices that replace or mimic the natural behaviour of the foot sole along with its sensory capabilities, for example in prosthetic applications, it is important to determine the sensing capabilities necessary for registering the full range of behaviourally relevant force patterns.

Here, our aim was to characterize the spatial properties and complexity of pressure patterns experienced on the foot sole during a range of natural everyday behaviour. We included tasks such as walking, running, jumping, and balancing, and recorded plantar pressure profiles in a young, healthy population wearing a standardized set of popular sports shoes. We analysed the locations and areas of contact on the foot sole, as well as the relationship between contact area and overall force. Finally, we quantified the complexity of spatiotemporal pressure patterns for different tasks, which yielded a small number of highly contacted independent regions whose locations suggest optimal sensor placements in future studies.

## Results

We recruited 20 young, healthy participants, who each executed a series of up to 15 different tasks spanning the range of everyday balancing and locomotion behaviours involving the foot sole (see Fig. 1d for illustrations and Methods for full details). These included balancing tasks, for example quiet standing, balancing on a wobble board, or rising from a sitting to a standing position, during which both feet were typically in contact with a surface simultaneously. We also included locomotion tasks, which comprised of walking at a number of speeds (slow, normal, fast, and jogging), up and down inclines and stairs, and on an uneven surface (gravel). In locomotion tasks typically only a single foot was on the ground at any given time. Finally, we included a number of jumping tasks, with participants instructed to jump from a low raised platform and land either on a single foot or on both feet. Participants wore a set of popular, standardized sports shoes in their size (Fig. 1a), which were outfitted with pressure sensitive insoles (Fig. 1b), capturing spatiotemporal pressure patterns with a spatial resolution of 3.9 sensors/cm^2^ (average of 622 active sensors per foot across participants) at a sample rate of 100 Hz. After calibration and pre-processing, the spatial pressure data were mapped onto a standardized foot outline (Fig. 1c) to allow localisation of sensors and joint analysis across participants (see Methods for details and Fig. 1e for example frames from different tasks).

**Fig. 1.**
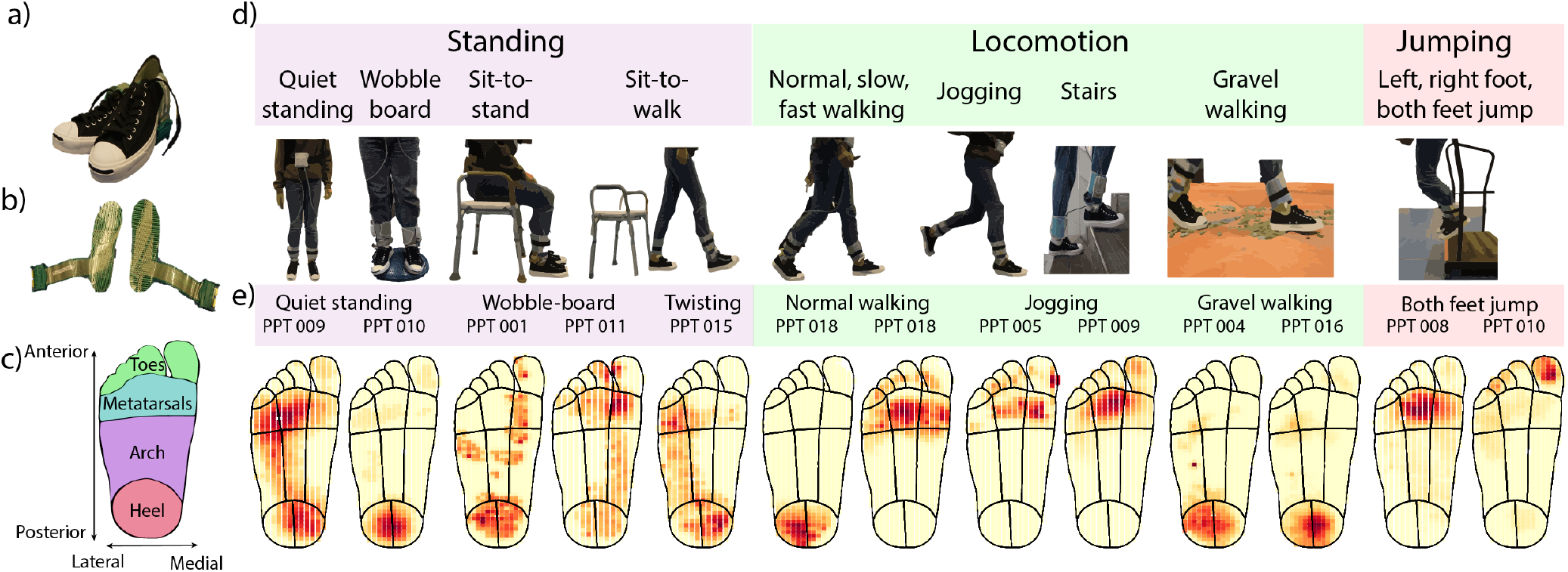
Overview of the setup, experimental tasks, and raw data. **a)** Standardized footwear worn by participants. **b)** TekScan®pressure insoles, recording pressure patterns with high spatial and temporal accuracy. **c)** Foot outline used for mapping, separated into the four coarse regions of the foot used during analysis. Anterior-posterior and medial-lateral axes labelled. **d)** The 15 tasks were split across three categories: standing, locomotion, and jumping. They included both stable and unstable surfaces and aimed to cover a range of everyday activities. **e)** Examples of individual frames from the pressure insole recordings.

We found that the average total force experienced by a single foot (expressed as percent body mass) was in agreement with previous studies across different tasks (7, 11, 12, 15): just below 50% body mass for standing tasks with both feet on the ground, and 75% body mass or higher for locomotion tasks, with overall forces increasing with speed and much greater than body mass during jogging (see Fig. S1 and Table S2). As expected, the highest forces were observed during jumping and jogging tasks, where loads exceeded body mass several-fold.

### Contact area

We first asked to what extent the foot sole was typically in contact with external surfaces and where on the foot sole contact was made. Because of the high spatial resolution of the pressure insoles, we were able to determine the contact area with high accuracy. During standing tasks, the average contact area experienced by the foot across all participants was around 58% of the foot, though this was highly variable both between participants and across the different tasks (Fig. 2a). During locomotion tasks, the average contact area was 48% of the foot. Interestingly, mean contact area remained similar across individual tasks within this category regardless of walking speed or surface. Indeed, while Kruskall-Wallis tests conducted across tasks within a given category were all significant (*p* < 0.020, see Table S7), many of the post-hoc pairwise Mann-Whitney U tests in the locomotion category were non-significant, primarily including walking on flat ground in comparison to jogging and walking on gravel (see Fig. 2a and Tables S8, S9, and S10). These findings are in direct contrast to the forces experienced in these tasks, which were highly significantly different (*p* < 0.001) within each task category (Table S3) as well as in all pairwise post-hoc comparisons (see Tables S4, S5, and S6). Contact area also varied considerably within a single task. Comparing the distribution of contact area over time across different tasks (Fig. 2b) confirmed the above findings: distributions differed between standing tasks, but were remarkably consistent and alsmost completely overlapping across locomotion tasks.

**Fig. 2.**
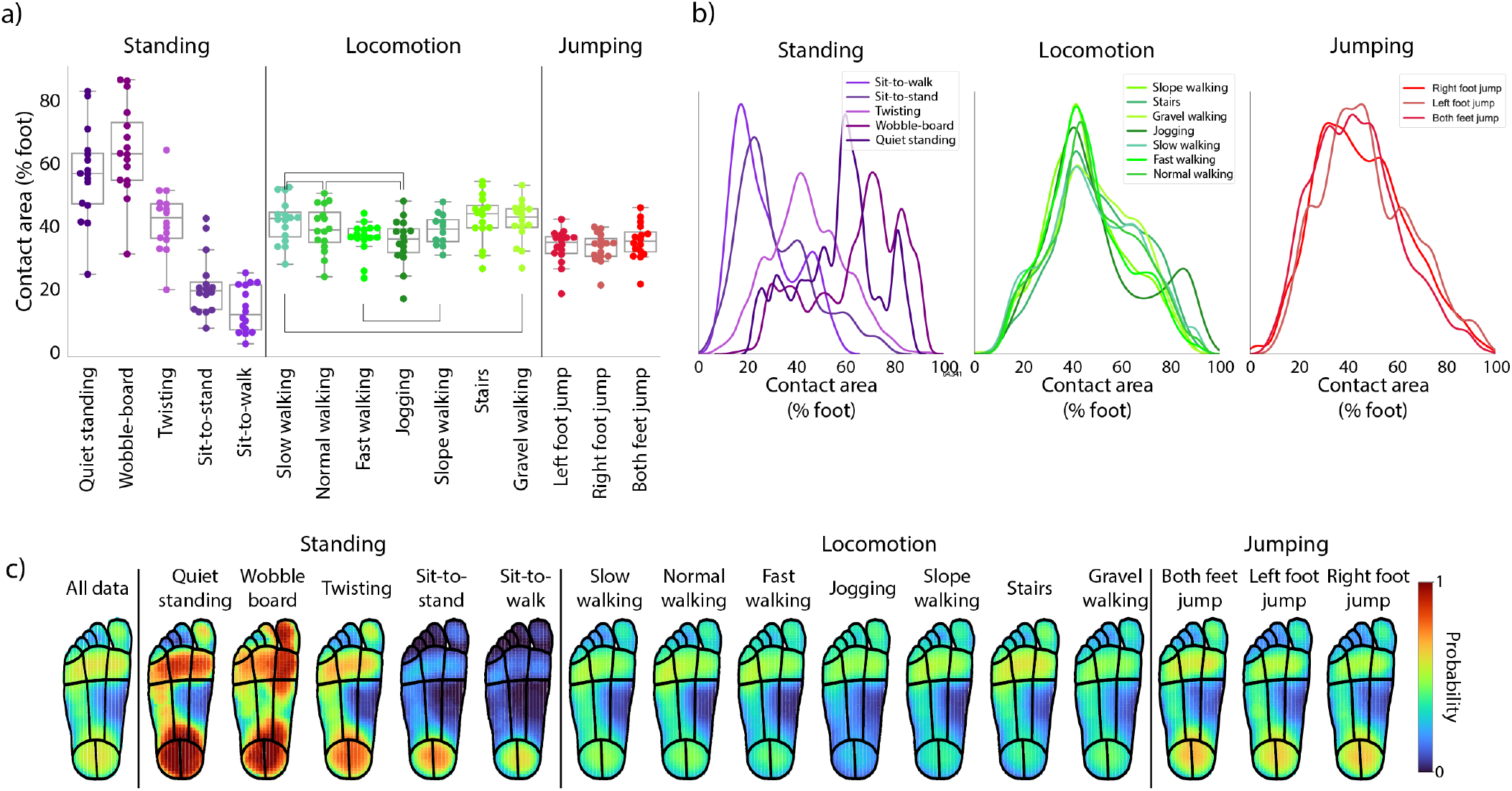
Contact area on the foot sole across tasks. **a)** Mean contact area experienced by either foot for all participants (coloured markers) across all tasks. Black horizontal lines indicate non-significant differences in Bonferroni-corrected post-hoc Mann-Whitney U tests. **b)** Distribution of contact area across individual tasks within each category. **c)** Probability of each sensor being in contact with the ground across all tasks, with red indicating high probability and blue indicating low probability.

Next, we quantified how often different regions of the foot made contact across the different tasks. As expected, contact probabilities were far from uniform. Overall, the heel and metatarsals were most likely to experience contact, followed by the lateral arch (see leftmost panel in Fig. 2c). During standing tasks, there was a clear difference between the areas of the foot likely to be in contact with the ground across individual tasks: when on the wobble-board, pressure was more likely to be spread evenly across the medial-lateral axis of the arch and the great toe was also more likely to be in contact with the ground relative to quiet standing. In the locomotion tasks, the pattern of contact areas across the foot was similar across the tasks, highlighting that the main regions involved in locomotion are the same irrespective of walking speed or surface.

### Contact area versus force

The analysis of contact area above suggested that the extent of contact of the foot sole might often not directly reflect the overall force applied by the foot. To directly test this relationship, we correlated force and contact area and calculated the proportion of shared variance for each task classification, including all tasks within each category. There was only a moderate relationship between force and contact area (*R*^2^ < 0.30 for all categories, see Fig. 3), suggesting that a greater area of the foot in contact with the ground does not necessarily mean a greater force experienced.

**Fig. 3.**
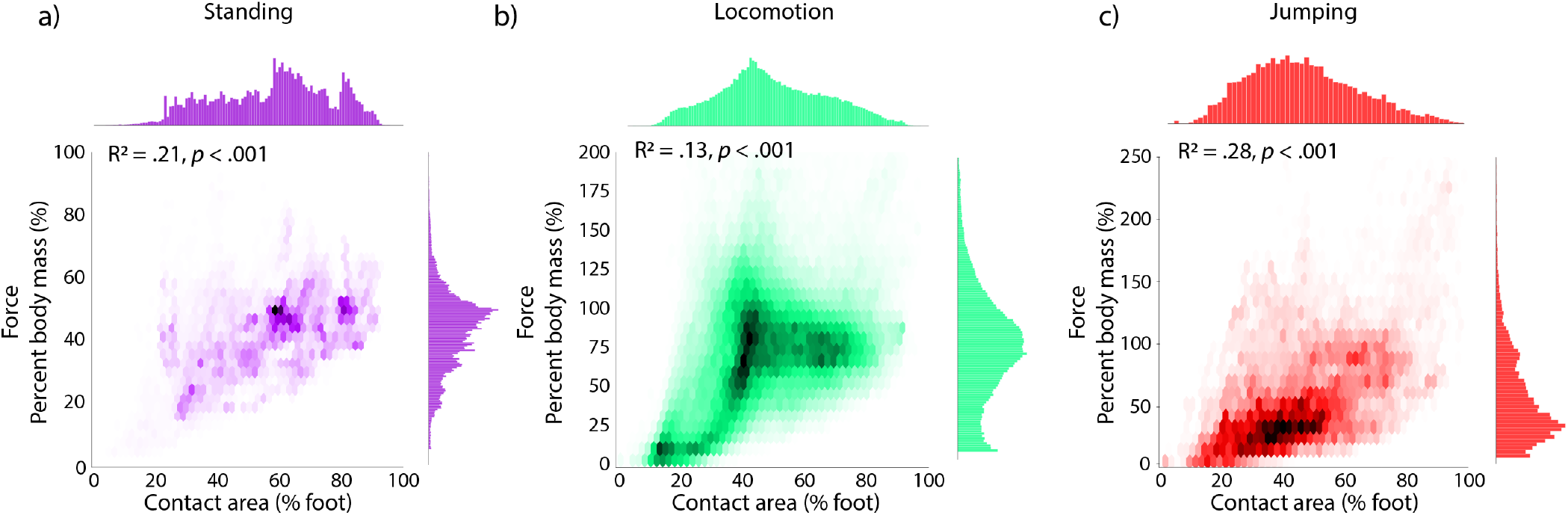
Relationship between force and contact area on the foot sole across task categories. 2D histograms showing relationship between overall force and contact area for **a)** standing, **b)** locomotion, and **c)** jumping categories, including data from all tasks within each category. Force and contact area exhibit a moderate positive correlation with there being a high density of data points clustered around 50% contact area and 75% body mass.

### Centre of pressure

The centre of pressure (CoP) on the foot sole is a popular measure in studies on gait and balance, mainly because of its relevance in mechanical models of balance (16). We investigated how often the CoP reflected the centre of a well-defined and relatively small contact region and how often it arose from a complex contact profile spanning larger areas of the foot. During standing tasks, the centre of pressure was primarily located at the arch (82% of time, see Fig. 4a), though with some variability between tasks (see example traces in Fig. 4c). Unlike quiet standing, CoP location was more evenly spread across the foot during locomotion and jumping (Fig. 4a). Contact area on the foot varied between CoP locations and was especially large for CoP locations on the arch, when on average 70% of the foot was in contact with the ground (Fig. 4b). These findings, combined with the fact that the probability that the arch was in contact with the ground was generally lower than that of the heel or metatarsals, suggested that often the foot might not actually make contact at the location where the CoP is located. Indeed, during standing tasks, the CoP was within the area of contact only 62% of the time (Fig. 4c), reducing to as little as 40% when standing on the wobble-board (see full break-down in Fig. S3). Conversely, the CoP was within the area of contact 92% of the time during locomotion tasks. When in an area of contact, the average pressure at the CoP was in the 74th percentile during standing and 90th percentile during locomotion, indicating that the CoP was not necessarily located where local pressure was highest. Overall, these results suggest that the CoP does not necessarily reflect the centre of single well-defined contact region, but, especially during balance, is a consequence of disparate contact regions spread across the foot sole.

**Fig. 4.**
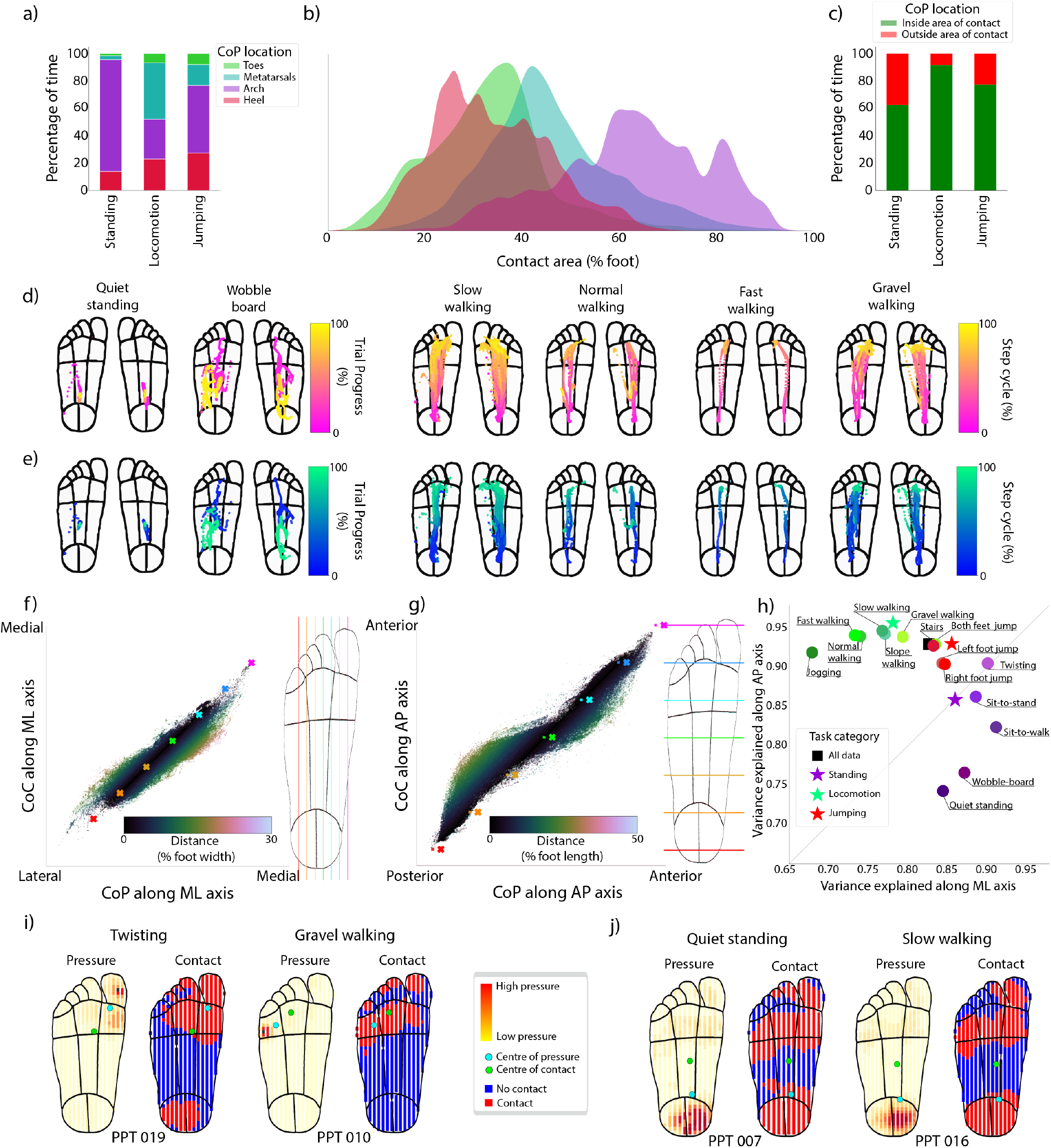
Centre of pressure characteristics and relationship between centre of pressure and centre of contact. **a)** Percentage of time spent by centre of pressure in different foot regions across task categories. **b)** Contact area when the CoP is in each of the foot regions across all tasks. **c)** Proportion of time the centre of pressure was in an area of contact for each task. **d)** Centre of pressure traces for one representative participant (PPT 004) across selected tasks. **e)** Centre of contact traces for same participant and tasks shown in c. **f)** Scatter plot showing CoP and CoC estimates along the medial-lateral axis of the foot (left). Coloured crosses indicate location along the medial-lateral axis off the foot as shown by vertical lines in the foot outline on the right. Each data point is shaded according to the distance between the CoP and CoC (see colourbar). **g)** Same as e, but for the anterior-posterior axis. **h)** Variance explained between the centre of pressure and centre of contact along the medial-lateral (horizontal) and anterior-posterior (vertical) axes across all tasks, task categories, and the whole data set. **i)** Examples of individual frames where the distance between the CoP and CoC is over 20% foot width. **j)** Examples of individual frames where the distance between the CoP and CoC is over 20% foot length.

To further investigate how the centre of pressure relates to the overall contact region, we also calculated the centre of contact (CoC), with each active sensor having an equal weight in the calculation (see Fig. 4c for CoP example traces and Fig. 4d for CoC example traces). Differences between the CoP and CoC suggest a skewed and possibly complex, rather than a flat distribution of the pressure across the contact area. Correlating the centres of pressure and contact, we noticed that along the medial-lateral axis, the CoP is typically located closer to the extremes of the foot than the CoC (Fig. 4e). Similarly, along the anterior-posterior axis of the foot, the CoP was also located closer to the extremes of the foot: more towards the heel than the CoC when considering the back half of the foot, but more towards the toes when considering the front half of the foot (Fig. 4f, see also examples in Fig. 4h). Overall, the shared variance between the CoP and the CoC was 83% along the medial-lateral axis and 90% along the anterior-posterior axis. While this indicates a strong relationship, there were occasions when the distance between the centre of pressure and centre of contact was over 25% of the foot width and 30% of the foot length. Interestingly, there were distinct differences in how well the CoP and the CoC agreed between task categories (Fig. 4g), with greater shared variance along the medial-lateral axis for standing tasks, while the opposite was true for locomotion tasks. This effect can be explained by the fact that during standing a large proportion of the foot (including the heel and metatarsals) was typically in contact with the ground at any time, but pressure was focused generally at one of these locations, biasing the centre of pressure in one direction along the AP axis while the centre of contact remains central on the foot, falling in an area of no or low contact. Conversely, during locomotion typically only a small region of the foot was in contact with the ground, especially during heel strike and toe off phases of the step cycle, and therefore there was less chance for the CoP to fall far outside of a localised contact area (see examples in Fig. 4h). Twisting and walking on gravel were the tasks during which the greatest distance between the CoP and CoC occurred, with the distance reaching up to 30% of foot width. This distance occurred due to localised pressure biasing the CoP, such as when stepping on a piece of gravel leading to a localised hotspot of pressure (Fig. 4h).

### Spatial complexity

Having established that pressure is often not uniformly distributed across the contact area and that the foot sole frequently makes contact at multiple locations, we set out to characterize the complexity of the pressure patterns in more detail. To do so, we used non-negative matrix factorization (NMF), a technique that decomposes high dimensional data sets (in this case spatial distributions of pressure recorded in each sampling frame) into a small set of components that reflect uncorrelated but commonly occurring pressure patterns. The number of components needed to reconstruct the pressure distributions over time with high accuracy indicates the complexity of the pressure patterns. Moreover, the location and extent of these components on the foot sole provide insight into which regions contribute frequently and independently to overall pressure patterns and suggest locations for efficient sensor placement in future studies. We chose NMF because it is similar to other popular dimensionality reduction techniques, such as principal component analysis, but forces both the components and their respective reconstruction weights to be positive, reflecting both the bounded nature of pressure patterns as well as potential neural mechanisms.

For each task, we used the number of components necessary to explain at least 90% of the variance in the pressure patterns as a measure of their complexity. On average, this number was 8 across all tasks, around 100 times fewer than the average number of active sensors, demonstrating that some of their signals were highly redundant. Nevertheless, there were clear differences in complexity across the different tasks (see Fig. 5a, d). Balancing on a wobble-board was more complex (11 components) than flat standing (4). Sit-to-stand and sit-to-walk required the fewest number of components (4 and 3, respectively). For locomotion tasks, walking on gravel was the most complex by far (16), reflecting the higher variability and more localised pressure peaks experienced in this task. There was little difference in the complexity between walking speeds (7, 6 and 7 for slow, normal and fast walking respectively), indicating that speed does not influence spatial complexity; instead, the environment (cf. gravel) appears to be the more important factor. These differences in complexity robustly emerged across different thresholds of variance explained (minimum threshold: 65%), with gravel walking and the wobble-board consistently requiring the highest number of components.

**Fig. 5.**
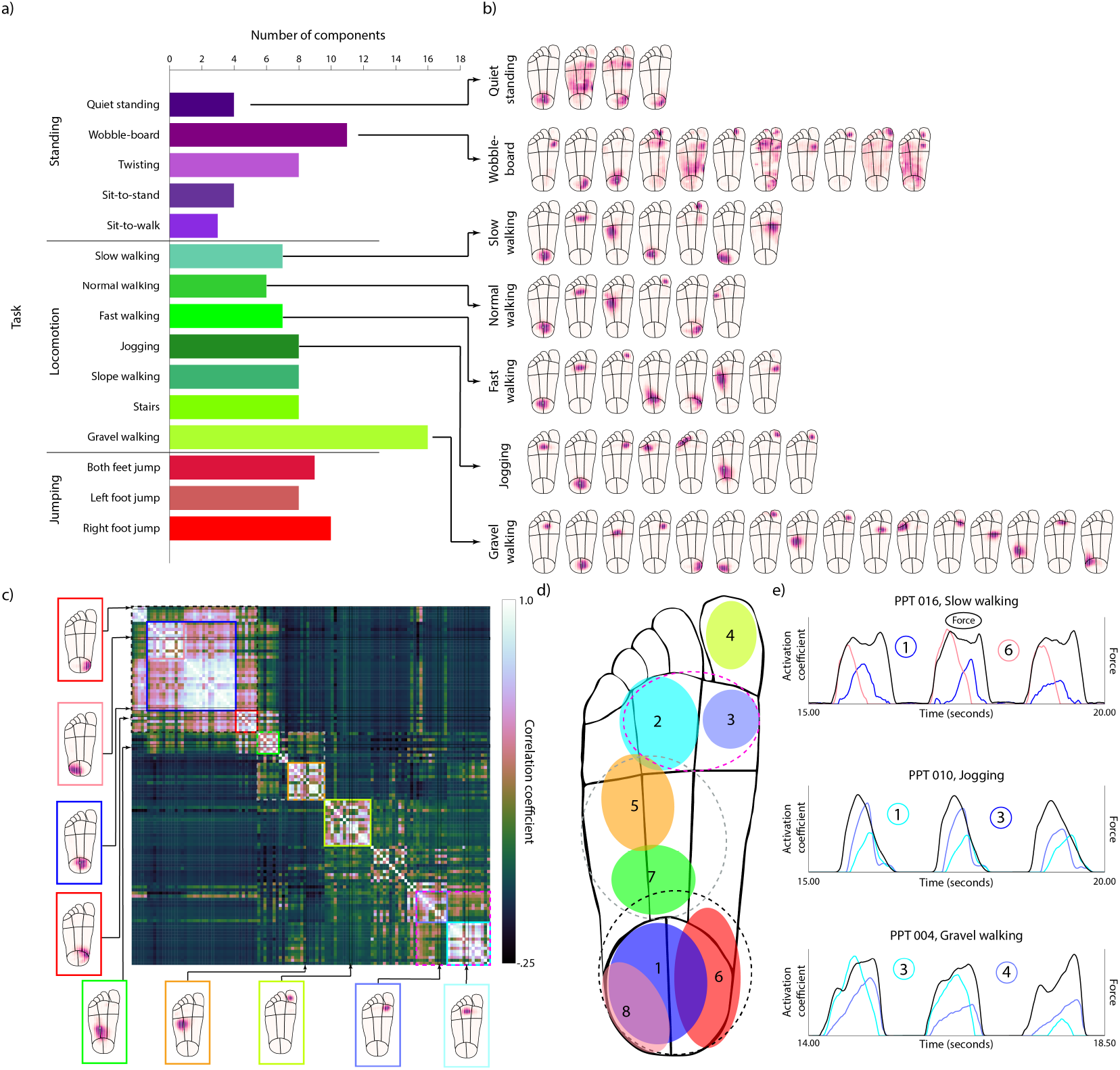
Spatial complexity of pressure patterns on the foot sole. **a)** Number of NMF components required to explain 90% of the variance in the spatial pressure distribution for different tasks. Many more components are required for the wobble-board and gravel walking tasks, demonstrating higher spatial complexity. **b)** Spatial plots of all components identified in panel a for selected standing and locomotion tasks, where darker shading indicates higher weight (normalized, arbitrary units). See Fig. S2b for components from all tasks. **c)** Pairwise correlations between all components required to explain 90% in each task, sorted to display clusters containing similar components along the diagonal. Coloured boxes with solid lines indicate distinct clusters that correspond to common regions on the foot. Boxes with dashed lines indicate overarching clusters that contain multiple distinct clusters. Examples of components that feature within each cluster are shown outside of the correlation matrix with arrows pointing to their respective row/column in the matrix. **d)** General locations (shown as coloured ellipses) of distinct clusters identified from the correlation analysis (panel c). Colours for clusters are the same as in c. Numbers indicate order in which clusters emerge, with smaller numbers indicating more prominent clusters. **e)** Examples of activation coefficients over time for selected NMF components and tasks. Line colour relates to cluster location in panel d and number relates to NMF component number in panel b.

Inspection of the individual NMF components revealed that components were generally well localised, covering only a small extent of the foot sole with typically a unimodal peak (see examples in Fig. 5b and the full set in Fig. S2b), suggesting that different localised regions make independent contributions to the overall pressure distribution. Some components related to standing tasks spanned a larger proportion of the foot and were sometimes multimodal, however their regions of maximum sensitivity were generally still relatively well localised.

The first two NMF components in most tasks were located on the heel and metatarsals, demonstrating the importance of these two regions specifically, and the anterior-posterior axis more generally. During more complex tasks, such when walking on gravel, twisting, and standing on a wobble-board, components were also located in regions that barely made any contact in more simple tasks and were much more localised, indicating much more varied pressure distributions in these tasks.

Interestingly, foot regions that are often considered as single functional units were often covered by multiple distinct components with small spatial offsets and the same configuration appeared reliably across different tasks. For example, for many tasks, multiple components were located across the metatarsals: the most prominent located centrally and second one located medially. Similarly, multiple components also emerged on the heel: a primary one on the central heel, flanked by medial and lateral components on either side. Importantly, these components did not simply reflect differential placement of the foot by participants, but instead could signal complex interactions with the ground over time even within single participants (see Fig. 5e for examples showing activation coefficients over time of nearby components located in the same foot region, which demonstrate temporal offsets between components and therefore a spatial shift in the pressure distributions).

Finally, to investigate which distinct foot regions emerged robustly across different tasks, we calculated pairwise correlations between the components derived from all tasks and then ran a cluster analysis (Fig. 5c). Eight distinct clusters were identified from the data, each responding to a unique location on the foot: the great toe, the medial and central metatarsals, the anterior and posterior lateral arch, and the medial, central, and lateral heel (see Fig. 5d for an illustration). These regions emerged in a robust order when increasing the overall number of components included in the cluster analysis: first the central heel, followed by the central and medial metatarsals, indicating their big contribution to pressure patterns across virtually all tasks. Next to be identified were the great toe, anterior lateral arch and medical heel, and then finally the lateral heel and posterior arch.

In summary, analysis of spatial complexity identified robust and localised components on the foot sole that explain most of the variance in spatial pressure pattern. Unstable or uneven ground increased the spatial complexity of pressure patterns considerably. Additionally, pressure patterns are divided on the foot more finely than often considered, specifically on the metatarsals and the heel, where multiple independent subregions contribute separately to overall pressure patterns.

## Discussion

In this study, we characterised the spatiotemporal pressure patterns that the plantar sole experiences during a range of common activities. During many of the tasks, slightly less than half of the foot sole was in contact with the ground on average. The resulting pressure distributions were not spatially uniform, but often skewed. Specifically, the centre of pressure was often biased by extreme, localised pressure. As a result, this measure, when taken alone, fails to capture subtleties in the stimuli experienced by the foot sole. Analysis of the spatial complexity of pressure signals revealed that the overall pressure distribution was well captured by a few localised components, mostly located at the heel and the metatarsals. These regions were each captured by multiple separate components, suggesting that specific, small subregions on the foot sole are differentially under load across different tasks and at different times.

### Selection of everyday tasks

To understand the role of tactile feedback during posture and gait, it is essential to first understand the stimuli that are experienced by the foot sole. Existing studies investigating pressure distributions and the role of tactile feedback typically involve only simple tasks, such as walking on flat surface and quiet standing (13, 14, 17, 18). While some research has begun to expand upon single-speed walking by investigating different speeds (19), we investigated a much broader range of tasks, including different walking speeds, surfaces and balance tasks of differing difficulty. The resulting data set allowed us to demonstrate how the distribution of pressure experienced by the foot sole differs greatly depending on the specific task, highlighting the importance of conducting a range of tasks to capture the full breadth in pressure patterns experienced during daily life.

### Identification of relevant foot regions

A major goal of the present study was to determine which regions of the foot sole are mainly involved in contact with the ground and make relevant and separable contributions to the overall pressure patterns. Previous work has typically divided the plantar sole into three or four regions: toes, metatarsals (sometimes merged with the toes to refer to the forefoot), arch (midfoot), and heel (rearfoot) (12, 18, 20). Often these mappings are inconsistent between studies, for example the rear foot can occupy up to 31% of foot length in some studies (12). When the foot sole is broken down further, there is no agreement on the number or extent of individual regions, with the arch sometimes being treated as a single region (21, 22) and other times as two regions split along the medial-lateral axis (13, 23). The lack of a common mapping technique between studies means limits generalisability and interpretation. Additionally, existing studies that have investigated optimal sensor placements have focused on locations that are recurrently active and allow for identification of heel strike and toe off through visual inspection (24), rather than sensors that provide information throughout a range of tasks.

Here, instead of pre-defining relevant foot regions, we used an unsupervised method where localised and relevant regions emerged from the data itself. By choosing a wide range of natural tasks, we ensured that the resulting findings were robust across multiple different tasks, including both walking and balancing. Eight independent spatial clusters emerged from this analysis, which captured most of the information contained in the pressure patterns across all tasks: the great toe, the medial, central and central metatarsals, the anterior and posterior lateral arch, and the medial, central and lateral heel. Interestingly, multiple components are located within regions that are often treated as singular, specifically the heel and the metatarsals. These results indicate a more complex interaction with the ground for these regions than commonly assumed, which might have different functional roles. It is an open question whether the tactile systems might interpret tactile feedback arising from these different sub-regions differently. On the engineering side, these results suggest that multiple sensors in these regions are necessary to capture a more accurate picture of the resulting pressure patterns.

Many clinical or research applications (for example neuroprosthetic devices) require pressure sensors to be placed across the foot sole to understand how the foot has been placed on the ground. While the pressure insoles used in the current study include hundreds of individual sensors, data efficiency, device robustness, and affordability generally preclude this option and only a very limited number of sensors is viable. Our findings suggest that in order to capture the maximum amount of information across daily tasks with a minimal amount of sensors, their locations should align with the clusters identified in the present study.

Finally, while the eight identified regions capture the majority of the complexity in the spatial pressure patterns, they do not capture all of it, and this is especially true for more challenging tasks, such as walking on gravel: in these tasks, localised pressure might arise in regions on the foot not included in our set (such as on the lateral metatarsals, see component 11 for gravel walking in Fig. S2b). It is possible that tactile feedback from these regions plays a functional role and indeed might be more relevant when tasks are difficult, so future research should study tasks that go beyond simple walking.

### Centre of pressure

The centre of pressure is often recorded as a measure of balance, with greater path lengths and variance of the CoP taken as an indication of poorer balance. However, as we have demonstrated, this metric can be biased by localised pressure when the ground is not flat, such as when walking on gravel. Additionally, the CoP is biased to the extremes of the foot, towards the toes and heels along the anterior-posterior axis, compared to the full area of contact, and this effect becomes more prominent with greater walking speed as the forces experienced at the heel and toes increase (25). Thus, information about pressure from regions close to the outer boundaries of the foot may be particularly relevant for the tactile system. Similarly, during quiet standing the CoP is mostly located at the arch, whereas most of the pressure signal is localised to the heel and metatarsals, while there may be little contact with the arch at all. This again suggests that any calculation of the centre of pressure by the tactile system must rely on sensory feedback from regions distant from the CoP and located towards the anterior and posterior boundaries of the foot. When the distance between CoP and CoC becomes greater, it can be identified that the environment is more complex, such as when walking on gravel. The tactile system must therefore be able to take into consideration both the location and magnitude of stimuli to keep track of the CoP in order to maintain balance and to help sense changes in the environment.

### Implications for tactile feedback processing

Tactile feedback from the foot sole contributes important information during walking and balance, as shown by increased sway and unsteady gait when this feedback is impaired (3–5). However, how exactly information about contact events is represented in tactile neural responses from the foot sole or how these are processed are currently open questions. Our analysis of spatial pressure patterns identified that at most 7, and often fewer, localized NMF components could explain 90% variance in pressure profiles in all tasks, with different components active at different times and during different tasks. This suggests that the tactile system might equally rely on a handful of localized feedback components, similar to the low-dimensional common set of muscle synergies in balance and walking (26). Feedback would also need to be integrated over large parts of the foot, for example when extracting the centre of pressure or similar measures relevant for balance. Finally, the large range of possible pressure values (from light contact to forces exceeding body weight several fold) necessitate neural mechanisms that are robust over this range. Recently, computational models have been developed that allow simulation of tactile neural responses from the foot and which will help study the nature of tactile feedback in more depth than possible at the moment (27).

### Limitations and future work

All participants in the current study were young, healthy adults. Pressure distributions change with age (12, 21, 28, 29), disease (22) and foot deformities (30, 31). Older adults walk slower than younger adults (5, 32, 33). The elderly also shift pressure in the anterior direction towards the toes (28) away from areas that exhibit the greatest loss in tactile sensitivity, with the forefoot experiencing the greatest pressure during adulthood and older adulthood compared with adolescence (12). Similarly, in diabetic patients with peripheral neuropathy, pressure sensitivity decreases across the entire foot (34) and peak pressure is greater (14, 35, 36), possibly to counter the increase in pressure sensitivity threshold observed in such populations (34). Changes in gait of patients with neuropathy are also exaggerated when walking on uneven surfaces (4, 37). Such alterations in gait observed in the elderly and neuropathic patients has been related to a decrease in sensitivity (14, 28, 38). These results demonstrate that sensitivity has a direct influence on plantar pressure experienced. The results of the current study are therefore not generalisable to populations in which changes in pressure are known to occur. Replication of the current study with older healthy and clinical populations will assist in the understanding of how tactile stimuli changes with age across a range of tasks.

The pressure patterns experience also depend on the footwear: the type of shoe worn influences how one places their foot on the floor and therefore resulting forces experienced (11, 18, 39, 40). The shoe itself, and specifically the thickness and stiffness of the sole, will also affect pressure patterns, with thicker soles spreading pressure more widely across the foot. In order to eliminate such effects in the present study, all participants wore the same brand and model of shoe. The shoe had a relatively thin sole, likely yielding more spatially fine-grained pressure patterns than might be one with different shoes. How exactly the shoe make affects pressure patterns will be an area of future investigation.

While using mobile in-shoe pressure measurement systems allows for much flexibility in task design and complexity, these devices come with inherent limitations. First, pressure sensitive insoles typically record lower forces than force plates (15, 41) and are harder to calibrate. Specifically, because each sensor only responds if its individual threshold is met, contact area (and pressure) will be underestimated when pressure is low and spread widely across a lot of sensors (42). However, the spatial resolution afforded by insoles is much greater than force plates which allowed for in depth analysis of spatial pressure patterns on a level finer than of individual regions of the foot. In the present study, high accuracy of the measured forces was not required, as we were mainly interested in spatial patterns. Second, performance decreases during high-impact activities, such as jogging (7). To minimise the effect of high-impact deterioration of insoles in the current study, jogging and jumping tasks were completed towards the end of the protocol and we measured insole performance both at the start and the very end of the protocol. Additionally, we restricted use of any individual insole pair to a maximum of three times before replacement. Whenever sensor dropout occurred during data collection, the pair would be replaced immediately. These actions ensured that insole performance was maintained as well as possible. Finally, the insoles used in this study are capable of recording normal force only. The foot sole also experiences shear forces, particularly during the heel strike and toe off phases of the step cycle (43), and some mechanoreceptors have been found to be sensitive to skin stretch (1). Insoles capable of measuring both pressure and shear have been recently developed (44) and will be useful for future research to arrive at a more complete picture of the complex force patterns experienced during natural activities.

## Methods

### Participants

20 participants (3 males and 17 females) with no history of gait irregularities or foot problems provided data for this study, with mean age 19.94 (standard deviation: 2.95) years old, mean height 169.74 (10.57) cm, and mean weight 64.73 (10.13) kg. 4 participants were excluded from data analysis, because their low weight prevented full calibration of the insoles. All participants provided informed consent prior to the start of data collection. The study protocol was approved by the ethical review board of the Department of Psychology at the University of Sheffield (protocol number 043136).

### Equipment and data collection

Participants wore the same kind of sports shoe (Converse Jack Purcell First In Class Low Top), available in UK sizes 2.5-12 (US Mens 3-12). For the current study, we used shoes between UK size 4 and 11 (modal participant size: 5). This shoe was chosen as it is commonly worn in daily life by all genders and features a relatively thin sole, thereby providing potentially rich tactile feedback.

Pressure sensitive insoles (TekScan®F-Scan®Sport Insoles, TekScan Inc., South Boston, Massachusetts, USA) were cut to each shoe size and inserted into both shoes. Each pressure sensor covered an area of 0.258 cm^2^, yielding around 650 sensors for UK shoe size 4 and up to around 1050 sensors for UK shoe size 11. Each sensor was sampled at 100Hz. The recording was started and finished during each task by the participants themselves by pressing a trigger linked to the pressure insoles. The pressure insole system connected via a local network to a recording laptop, allowing the participant to move unrestricted.

Participants also wore three Opal^TM^ wireless inertial measurement units (IMUs; APDM Wearable Technologies, Portland, Oregon, United States) throughout the experiment. Two IMUs were placed on the ventral side of both ankles and one was placed on the lumbar spine (L5). All IMUs were attached directly to the skin using double sided medical tape and secured with microporous tape. Signals from the IMUs were used during data segmentation, for example to identify when participants were walking versus turning.

### Sensor calibration and validation

To calibrate the pressure insoles, participants first performed a step calibration protocol, provided by the TekScan research software, during which participants stood on one leg, before transitioning to the other leg. Step calibration fits the initial response of the insoles to the load applied by the participant, and also estimates and corrects for the sensor drift over time. This calibration can fail for participants with low weight, causing four participants to be excluded from the sample. Visual inspection of the live recordings in the TekScan software was used to identify any major issues in data collection. One further calibration trial was then conducted to identify the outer boundaries of the foot outline. To do this, participants were asked to roll their foot over a ball trying to cover the whole foot. Next, four validation trials were conducted to measure insole performance: 1) participants initially stood on two feet before shifting to one foot at a time and maintaining each posture for 10 seconds, 2) participants began with two feet flat on the ground, before shifting to standing on their heels and toes, each for 10 seconds, 3) participants began on two feet before shifting to the left foot and 4) participants began on two feet before shifting to the right foot. Regardless of the number of feet on the ground or posture, the total force applied during static stance should correspond to the participant’s mass. The first validation trial was repeated at the start and end of the study to allow measurement of insole performance over the study. Expected participant mass during single foot standing was calculated. Based on the average calculated expected mass over a 5 second time period during the first validation trial, a constant was generated to recalibrate the pressure data during pre-processing. Following recalibration, the average participant mass during this 5 second time period would equal to the true participant mass.

Compared to standing flat on a single foot (against which the insoles were calibrated), we noted underestimation of the total force when both feet were flat on the ground by on average 10% and an overestimation when standing on a single foot by 25% during heel stance and 21% during tiptoe stance. These effects were likely due to individual sensors not reaching threshold when the overall load was spread between many sensors. Insole performance also decreased over time: the force recorded during single foot stance decreased by an average of 20% for either foot during the course of the experiment.

### Experimental tasks

Participants executed up to 15 individual tasks (see Table 1 for list and Table S1 for full details), designed to encompass a wide range of activities that occur in daily life. These tasks belonged to one of three different categories: standing, locomotion and jumping. Three participants were unable to complete slope walking, which was conducted outside, due to adverse weather at the time.

**Table 1.** All experimental tasks carried out by the participants, grouped by task category. Full task instructions are in Table S1.

| Category | Task |
| --- | --- |
| Standing | Quiet standing<br>Standing on a wobble board<br>Sit-to-stand<br>Sit-to-walk<br>Standing twists |
| Locomotion | Normal speed<br>Fast<br>Slow<br>Up/down a slope<br>Up/down stairs<br>Jogging<br>Walking on gravel |
| Jumping | Drop jump onto both feet<br>Drop jump onto the left foot<br>Drop jump onto the right foot |

### Data preprocessing

All data processing and analysis was carried out in Python (version 3.8.5) using NumPy (version 1.19.1), Pandas (version 3.8.2), Scipy (version 1.5.0) and Scitkit-learn (version 1.1.1).

#### Filtering

To filter out sensor noise, an isotropic Gaussian filter with standard deviation 0.5 (in sensor space) was applied to the spatial dimension of the data, and a Butterworth filter with a frequency cut-off of 18 Hz was applied to the temporal dimension.

#### Mapping the pressure data onto a standardized foot

To generate a standardized foot with accurate proportions, the length and width of participants’ heel, arch, metatarsals and great toe were measured as a proportion of overall foot width and length. The length and width of each region was averaged to generate the standardized foot outline. To map the data onto the foot, the filtered pressure data from all tasks was first concatenated. Then, the outer borders of the pressure matrix along both the anterior-posterior and medial-lateral axes were determined by identifying rows or columns in the sensor matrix that contained non-negative pressure values across any of the tasks, and empty matrix rows and columns were removed. Within the model of the foot outline, a matrix was created with a spacing corresponding to that of the pressure insole (see Fig. S4b). Each recorded pressure value was then mapped onto its corresponding location within the foot outline. On average, this mapping captured around 622 sensors (range: 459 to 826), and on average 94% of the total recorded pressure (see Fig. S4d for mappings for all participants).

#### Aligning the insole and IMU data

Once recording was started for each task, participants were asked to stomp one foot, which led to easily identifiable spikes in both the pressure data and the IMU signals, which were used to synchronize both data streams. The stomp itself was deleted from the data and its time was used to signal the onset of the task.

#### Data trimming for standing tasks

For data analysis, data was trimmed to only include relevant aspects of each task. For quiet standing the initial 10 seconds after task onset were cut and the next 45 seconds were used in the analysis. This enabled participants to settle into a comfortable posture. 35 seconds of time stood on the wobble-board was used for analysis, beginning 10 seconds after the stomp. This was to ensure that participants were able to mount the board and find their balance. Sit-to-stand and sit-to-walk trials were designed to measure the forces during the process of getting up to a standing position. Therefore, data from sit-to-walk was analysed up until the participant began to step as identified using manual segmentation of the pressure data.

#### Identifying steps within the pressure data

Within all locomotion tasks, individual steps were extracted by identifying periods where the pressure signal was below a given threshold. This threshold was typically set to 500 kPa experienced across the entire foot, but was adjusted to 3000 kPa and 5000 kPa for participants 14 and 16 respectively, due to excessive contact between foot and insole when the foot was off the ground. Turns during walking were identified automatically when the gyroscope within the IMU registered a rotation of at least 45°and a turn angle of at least 115°. Manual checks and, if necessary, adjustments were made to ensure that turns were correctly identified: if any part of a step occured while turning, as identified using the IMU placed on the lower back, or after the number of stairs climbed was completed, the step was ignored. The first and last step were removed before normalizing the length of all steps to 100 time points.

### Data analysis

#### Force and contact area

Total force was calculated by summing the signals across all sensors on a given foot and expressed as a percentage of the participant’s body mass. Contact area was calculated based on the proportion of active pressure sensors within the standardized foot outline. Timepoints during which total force values under 5% body mass were excluded from further analysis as these likely referred to noise caused by static contact between the insole and the foot when not on the ground. To examine differences in force and contact area between tasks within a category, Kruskall-Wallis tests were conducted, as all data violated the assumptions of normality and equality of variance. Bonferroni corrected post-hoc Mann-Whitney U tests were conducted to identify significant pairwise comparisons within task categories. To investigate the relationship between force and contact area, a Spearman’s Rho correlation was conducted.

#### Centre of pressure and centre of contact

Centre of pressure was calculated using the weighted average of all active pressure sensors. Centre of contact was calculated with each active sensor contributing an equal weight to the average.

#### Spatial complexity of pressure patterns

Non-negative matrix factorisation, a dimensionality reduction technique, was implemented along the spatial dimension of the data. Insole data from all participants was first scaled to a common size (UK size 11) and interpolated to fit the sensor grid for this show size, prior to mapping onto the standardized foot, ensuring that the data from all participants was of the same dimensionality: a size 11 foot with matrix dimensions of 56×20. NMF components are learned simultaneously and can therefore differ depending on how many components are extracted. To test whether the number of components affected the results, NMF models were calculated with one to thirty components, with the optimization run ten times each. Results of these simulations were consistent for each iteration, allowing for final analysis to be run once per number of components (1–30). To identify regions on the foot which were frequently occupied by sensors with high weights in individual NMF components across tasks, we calculated pairwise correlations between all components that were required to explain 90% variance in each individual task. The resulting correlation matrix was then sorted to identify clusters by calculating the distance between component pairs using Scipy’s clustering package.

## ACKNOWLEDGEMENTS

LDC was supported by a studentship from the MRC Discovery Medicine North (DiMeN) Doctoral Training Partnership (MR/N013840/1). HMR was supported by a studentship from the EPSRC Doctoral Training Partnership (EP/T517835/1). We would like to thank Miguel Casal for advice on the NMF analysis. This manuscript was typeset using the Henriques Lab Overleaf bioRxiv template available at https://henriqueslab.github.io/resources/bioRxivTemplate/.

## AUTHOR CONTRIBUTIONS

LDC designed the protocol with guidance from CM and HPS. LDC and HMR conducted data collection. LDC conducted data analysis, with guidance from HPS. Figures were generated by LDC, with feedback provided by all authors. LDC and HPS wrote the manuscript. All authors reviewed and approved the manuscript.

## COMPETING FINANCIAL INTERESTS

The authors declare no conflicts of interest.

## DATA AVAILABILITY

All raw data and code used to process and analyse the data is available at *https://osf.io/n9f8w/*, DOI: 10.17605/OSF.IO/N9F8W.

## Supplementary material

**Table S1.**
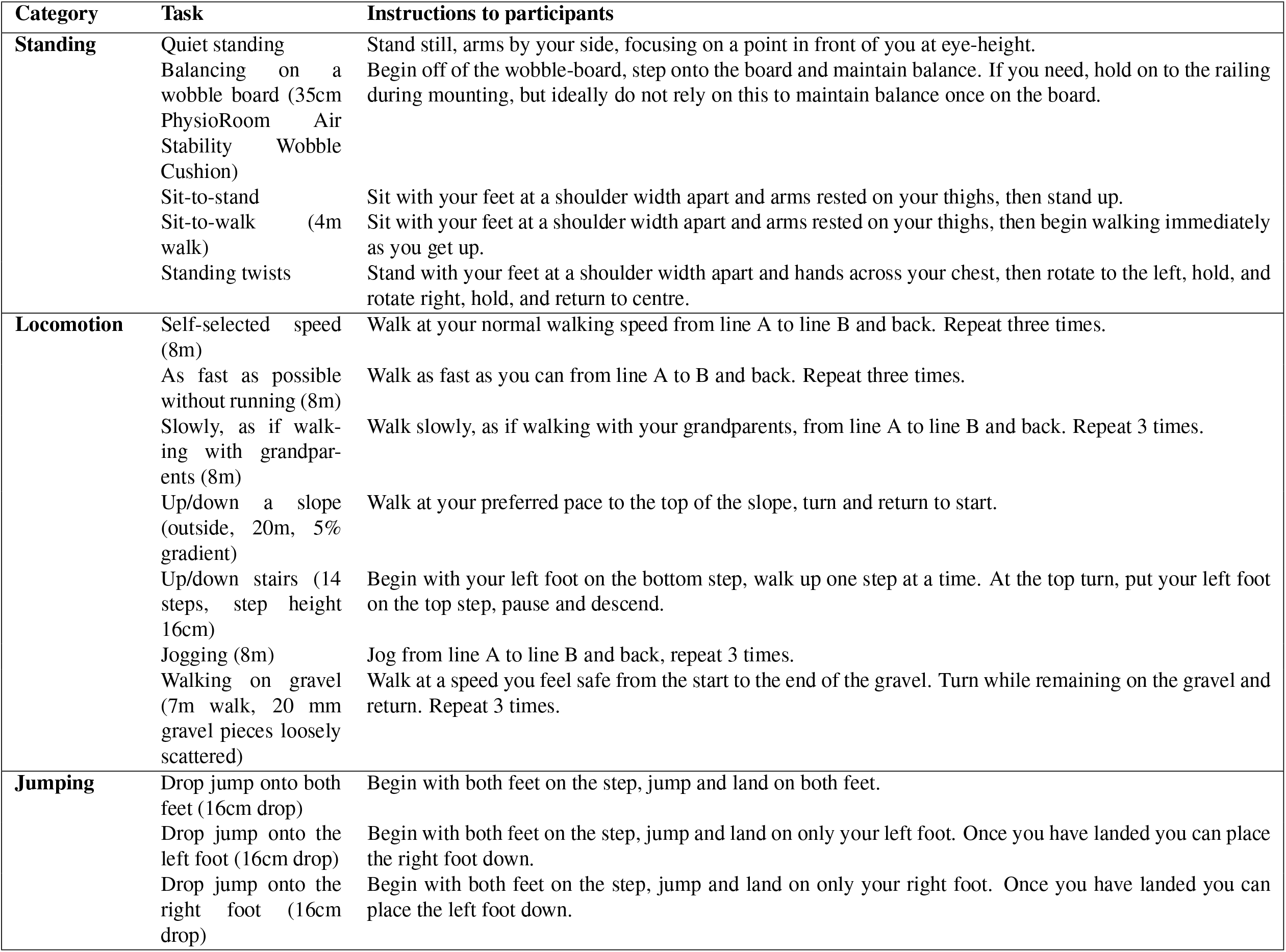
All tasks carried out by the participants, grouped by task category.

**Table S2.**
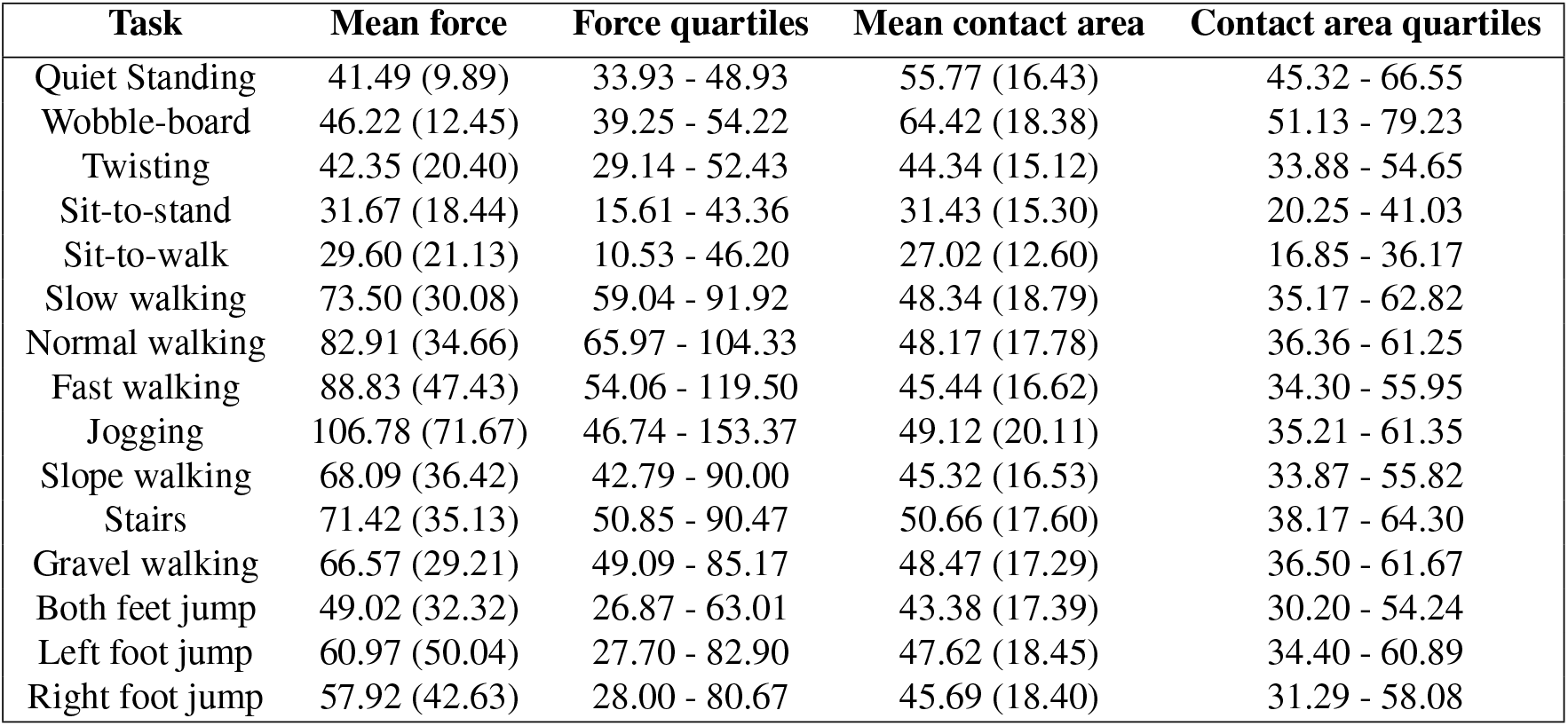
Mean (SD) of pressure and contact area across tasks averaged over participants and interquartile ranges.

| Task | Mean force | Force quartiles | Mean contact area | Contact area quartiles |
| --- | --- | --- | --- | --- |
| Quiet Standing | 41.49 (9.89) | 33.93 - 48.93 | 55.77 (16.43) | 45.32 - 66.55 |
| Wobble-board | 46.22 (12.45) | 39.25 - 54.22 | 64.42 (18.38) | 51.13 - 79.23 |
| Twisting | 42.35 (20.40) | 29.14 - 52.43 | 44.34 (15.12) | 33.88 - 54.65 |
| Sit-to-stand | 31.67 (18.44) | 15.61 - 43.36 | 31.43 (15.30) | 20.25 - 41.03 |
| Sit-to-walk | 29.60 (21.13) | 10.53 - 46.20 | 27.02 (12.60) | 16.85 - 36.17 |
| Slow walking | 73.50 (30.08) | 59.04 - 91.92 | 48.34 (18.79) | 35.17 - 62.82 |
| Normal walking | 82.91 (34.66) | 65.97 - 104.33 | 48.17 (17.78) | 36.36 - 61.25 |
| Fast walking | 88.83 (47.43) | 54.06 - 119.50 | 45.44 (16.62) | 34.30 - 55.95 |
| Jogging | 106.78 (71.67) | 46.74 - 153.37 | 49.12 (20.11) | 35.21 - 61.35 |
| Slope walking | 68.09 (36.42) | 42.79 - 90.00 | 45.32 (16.53) | 33.87 - 55.82 |
| Stairs | 71.42 (35.13) | 50.85 - 90.47 | 50.66 (17.60) | 38.17 - 64.30 |
| Gravel walking | 66.57 (29.21) | 49.09 - 85.17 | 48.47 (17.29) | 36.50 - 61.67 |
| Both feet jump | 49.02 (32.32) | 26.87 - 63.01 | 43.38 (17.39) | 30.20 - 54.24 |
| Left foot jump | 60.97 (50.04) | 27.70 - 82.90 | 47.62 (18.45) | 34.40 - 60.89 |
| Right foot jump | 57.92 (42.63) | 28.00 - 80.67 | 45.69 (18.40) | 31.29 - 58.08 |

**Table S3.** Results from Kruskall-Wallis test run on forces per task category. * indicates significance level of p<.001.

| Task category | Levene's test of homogeneity of variance | Shapiro-Wilk test of normality | Kruskal-Wallis H |
| --- | --- | --- | --- |
| Standing | 5585.14* | 0.98* | 12649.44* |
| Locomotion | 4499.14* | 0.95* | 9102.16* |
| Jumping | 280.47* | 0.83* | 230.26* |

**Table S4.** Bonferroni corrected post-hoc Mann-Whitney U tests run on forces in each task within the standing task category. U statistic shown for each pairwise comparison. * indicates significance level of p<.008.

| Task | Wobble-board | Twisting | Sit-to-stand | Sit-to-walk |
| --- | --- | --- | --- | --- |
| Quiet standing | 5246265152* | 2194534021* | 319399852* | 118829568* |
| Wobble-board | - | 1688180535* | 231962102* | 85485649* |
| Twisting | - | - | 59971375* | 23037320* |
| Sit-to-stand | - | - | - | 2060881* |

**Table S5.**
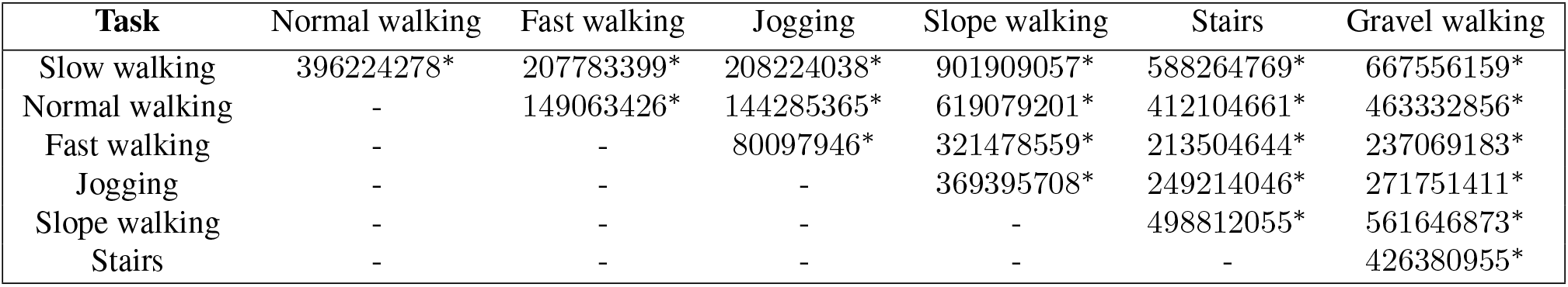
Bonferroni corrected post-hoc Mann-Whitney U tests run on forces in each task within the locomotion task category. U statistic shown for each pairwise comparison. * indicates significance level of p<.002.

**Table S6.** Bonferroni corrected post-hoc Mann-Whitney U tests run on forces in each task within the jumping task category. U statistic shown for each pairwise comparison. * indicates significance level of p<.017.

| Task | Left foot jump | Right foot jump |
| --- | --- | --- |
| Both feet jump | 47581465* | 43648768* |
| Left foot jump | - | 58287796 |

**Table S7.** Results from Kruskall-Wallis test run on contact area per task category. * indicates significance level of p<.001.

| Task category | Levene's test of homogeneity of variance | Shapiro-Wilk test of normality | Kruskal-Wallis H |
| --- | --- | --- | --- |
| Standing | 564.15* | 0.98* | 37813.56.88* |
| Locomotion | 249.03* | 0.99* | 1962.17* |
| Jumping | 25.77* | 0.98* | 240.93* |

**Table S8.** Bonferroni corrected post-hoc Mann-Whitney U tests run on contact area in each task within the standing task category. U statistic shown for each pairwise comparison. * indicates significance level of p<.008.

| Task | Wobble-board | Twisting | Sit-to-stand | Sit-to-walk |
| --- | --- | --- | --- | --- |
| Quiet standing | 4954841944.5* | 2950107770* | 399221907.5* | 156986859* |
| Wobble-board | - | 2191026239* | 278485944.5* | 107036012* |
| Twisting | - | - | 68137860.5* | 27565498.5* |
| Sit-to-stand | - | - | - | 2232182.5* |

**Table S9.** Bonferroni corrected post-hoc Mann-Whitney U tests run on contact area in each task within the locomotion task category. U statistic shown for each pairwise comparison. * indicates significance level of p<.002.

| Task | Normal walking | Fast walking | Jogging | Slope walking | Stairs | Gravel walking |
| --- | --- | --- | --- | --- | --- | --- |
| Slow walking | 480922411 | 275225609.5* | 281561941.5 | 857467504.5* | 500137809.5* | 561420207 |
| Normal walking | - | 170775152.5* | 174941063.5 | 532407301.5* | 307808629.5* | 345618210.5* |
| Fast walking | - | - | 84086896.5* | 255914915.5 | 144803053.5* | 163786346.5* |
| Jogging | - | - | - | 305441970.5* | 178954122.5* | 200041536.5* |
| Slope walking | - | - | - | - | 445064635.5* | 504008676* |
| Stairs | - | - | - | - | - | 421288383* |

**Table S10.** Bonferroni corrected post-hoc Mann-Whitney U tests run on contact area in each task within the jumping task category. U statistic shown for each pairwise comparison. * indicates significance level of p<.017.

| Task | Left foot jump | Right foot jump |
| --- | --- | --- |
| Both feet jump | 46827200* | 45351193.5* |
| Left foot jump | - | 60974979* |

**Fig. S1.**
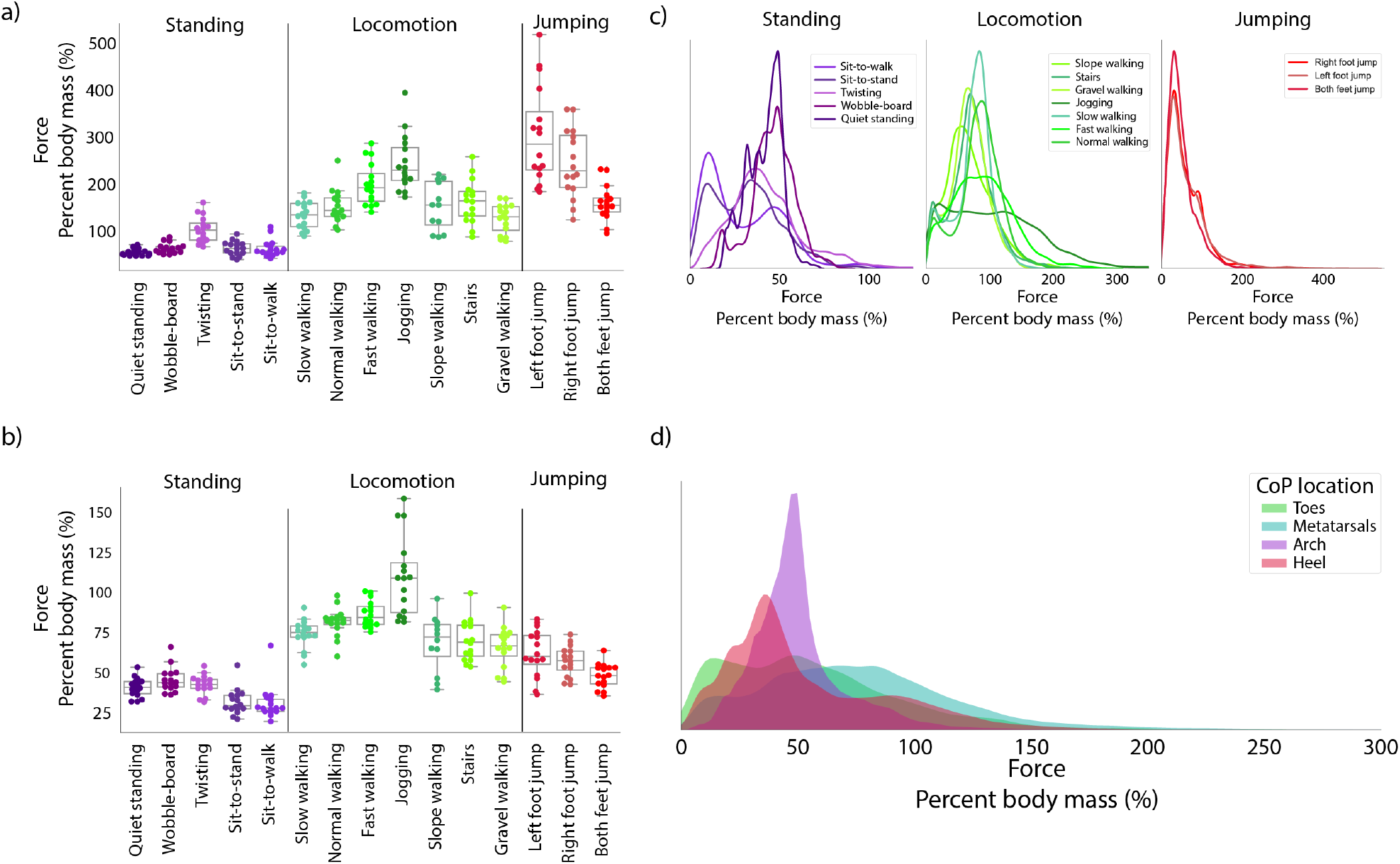
Force experienced by the foot across different tasks. **a)** Maximum force experienced by either foot for all participants (coloured markers) across all tasks. Boxplots show the distribution of maximum force over participants. **b)** Mean Force per task. **c)** Distribution of force experienced across each task within categories. **d)** Force when the CoP is in each of the foot regions across all tasks.

**Fig. S2.**
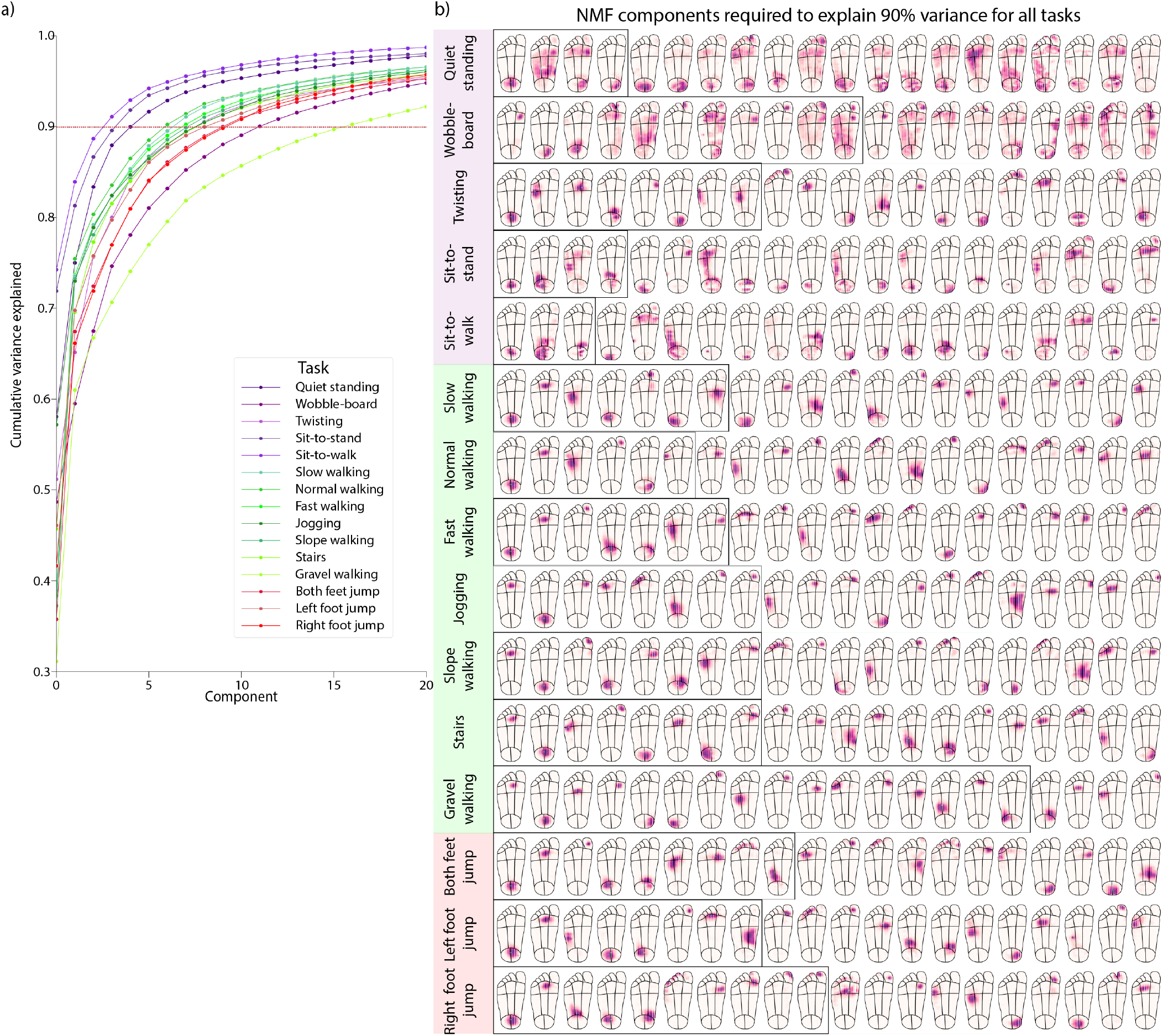
**a)** Scree plot showing variance explained by each component across tasks. **b)** The first 20 NMF components for all tasks plotted spatially on the foot. Darker purple indicates higher weight. The black box indicates the components required to explain 90% variance in the data for that tasks.

**Fig. S3.**
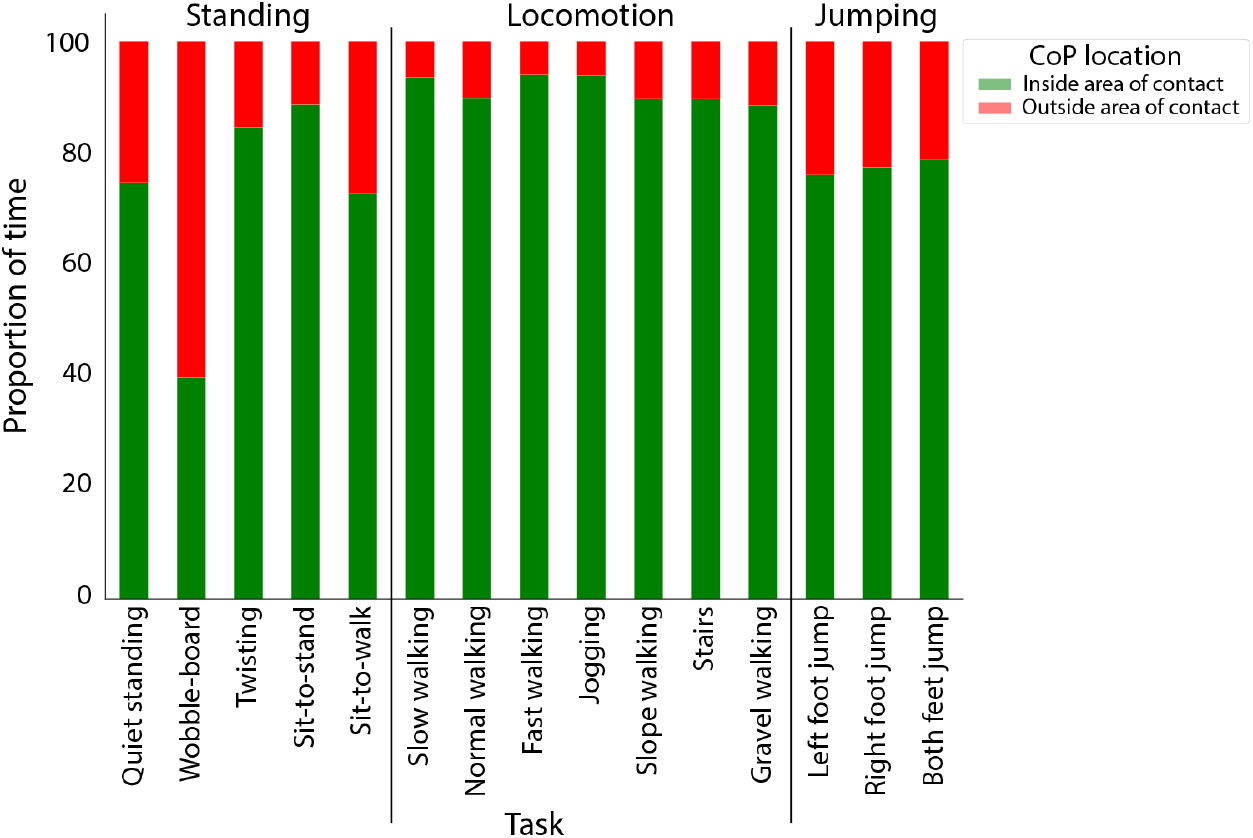
Proportion of time the centre of pressure is within an area of contact for each task.

**Fig. S4.**
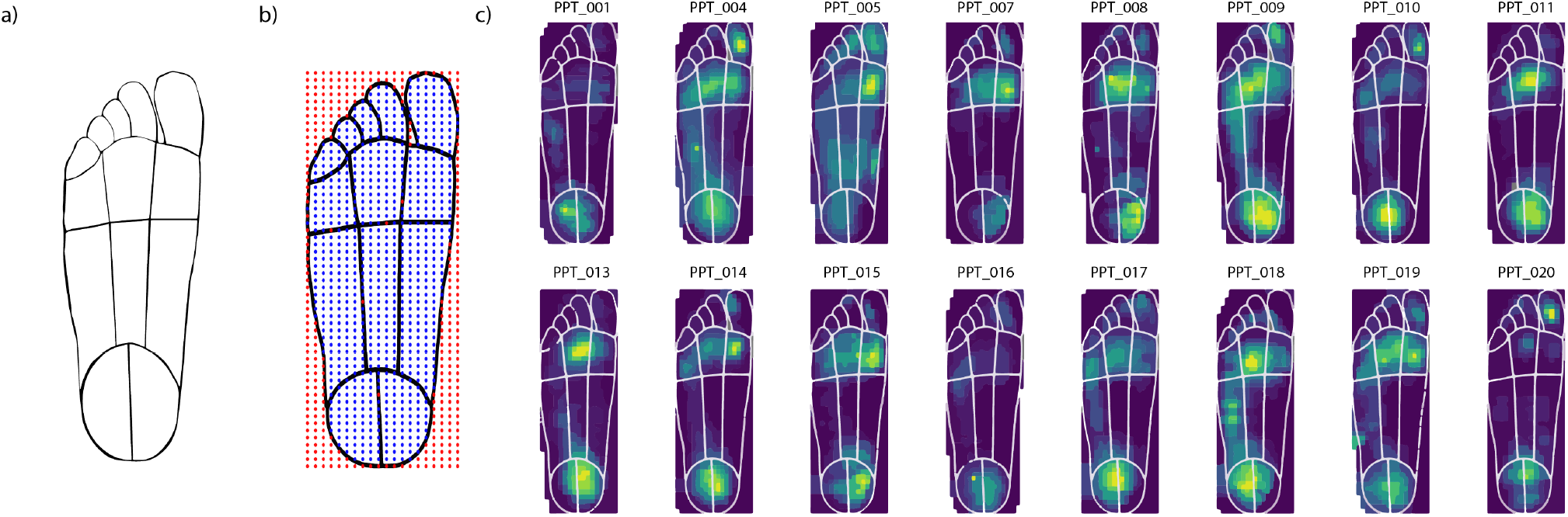
**a)** Standardized foot outline used to map the pressure data onto. **b)** Sensors mapped onto the foot outline, where blue points are sensors mapped onto the foot and red points reflect (potential) sensors not on the foot. **c)** Demonstration of how the raw pressure data aligns with the standardized foot. Mean pressure across all tasks for each participant with the standardized foot outline overlaid. Yellow indicates higher pressure, purple indicates low pressure.

## Notes

https://osf.io/n9f8w/

### Competing Interest Statement

The authors have declared no competing interest.

